# Risk practices for bovine tuberculosis transmission to cattle and livestock farming communities living at wildlife-livestock-human interface in northern KwaZulu Natal, South Africa

**DOI:** 10.1101/699520

**Authors:** Petronillah Rudo Sichewo, Catiane Vander Kelen, Séverine Thys, Anita Luise Michel

## Abstract

Bovine tuberculosis (bTB) is a disease of cattle that is transmitted through direct contact with an infected animal or ingestion of contaminated food or water. This study seeks to explore the local knowledge on the disease and establish the risk practices that lead to its transmission to cattle and humans (zoonotic TB) in a traditional livestock farming community with a history of bTB diagnosis in cattle and wildlife. Information was collected using a qualitative approach of Focus Group Discussions (FGDs) targeting household members of livestock farmers that owned either bTB infected or uninfected herds. We conducted fourteen FGDs (150 individuals) across four dip tanks that included the following categories of participants from cattle owning households: heads of households, cattle keepers, dip tank committee members and women. The qualitative data was managed using NVivo Version 12 Pro^®^software. Social and cultural practices were identified as major risky practices for bTB transmission to people, such as the consumption of undercooked meat, consumption of soured /raw milk and lack of protective measures during slaughtering of cattle. The acceptance of animals into a herd without bTB pre-movement testing following traditional practices (e.g. *lobola*, ‘bride price’, the temporary introduction of a bull for ‘breeding’), the sharing of grazing and watering points amongst the herds and with wildlife were identified as risky practices for bTB transmission to cattle. Overall, knowledge of bTB in cattle and modes of transmission to people and livestock was found to be high. However, the community was still involved in risky practices that expose people and cattle to bovine TB. An inter-disciplinary ‘One Health’ approach that engages the community is recommended, to provide locally relevant interventions that allows the community to keep their traditional practices and socio-economic systems whilst avoiding disease transmission to cattle and people.

**Author summary:** Bovine tuberculosis (bTB) is a respiratory disease of cattle that is transmitted to other animals as well as humans (zoonotic TB) through direct contact with infected animals, and consumption of contaminated food (animal products) or water. The study explains the complexities of human-animal relations, reflects on how people understand and conceptualize risk of bovine tuberculosis (bTB) in an endemic area considering the economic value of livestock keeping as well as social and cultural practices of importance to the community. The results of this study identified socio-cultural practices that involved consumption of raw or undercooked animal products and handling of infected animal products during animal slaughter as major risky practices for bTB transmission to people. Introduction of animals into a herd without bTB testing for socio-cultural purposes and sharing of resources amongst the communal herd and with wildlife were identified as risky practices for bTB transmission to cattle. The findings of this study illustrate the need for a One Health strategy that develops appropriate public health policy and related education campaigns for the community as control of zoonotic TB in people depends on the successful control of bovine TB in cattle.

## Introduction

Bovine tuberculosis (bTB) is chiefly a chronic respiratory disease of cattle caused by the bacteria *Mycobacterium bovis* (*M. bovis*) that has multiple incidental hosts including humans, goats, cats, dogs and wild animals [1–3]. In South Africa, the disease is a state controlled disease in cattle due to its negative impact on livestock production, export and local market in animal and animal products, wildlife conservation efforts and could increase human health costs [4,5]. Similar to most African countries where bTB is prevalent, the surveillance and ‘test and slaughter’ programs are not optimally implemented in South Africa due to a lack of resources, despite the potential hazard to human health [6,7]. *M. bovis* infection in humans is referred to as zoonotic tuberculosis and has been classified by the World Health Organisation as a neglected zoonotic disease [8]. Zoonotic TB transmission to humans is predominantly through the consumption of contaminated animal products such as unpasteurised dairy products and less frequently attributed to animal-to-human or human-to-human through direct contact [9].

The study area is surrounded by conservation areas where *M. bovis* infection has been established in African buffalo which is wildlife maintenance host [10]. Recent studies have revealed *M. bovis* infection in communal cattle in the same area with 28% of the farmers having at least one test positive animal in their herd [11].The presence of *M. bovis* in cattle and wildlife increases the risk of zoonotic TB transmission to susceptible human populations living at the wildlife/livestock/human interface [5,12]. A lack of or insufficient implementation of the ‘test and slaughter’ disease control scheme, consumption of uncooked meat products and soured milk, poor understanding of zoonotic TB and poor sanitary conditions are some of the potential risk factors for *M. bovis* infection and disease in humans [13].

Due to an increase in land use there is infringement of human activities into conservation areas that results in the sharing of natural resources with wildlife [14]. Studies in other countries have suggested that transmission of bTB to cattle from wildlife occurs either through direct contact at shared resources such as watering points or indirectly, when cattle graze on contaminated pastures [15]. Therefore, the wildlife-livestock interface has been defined as a high-risk area for bTB transmission from wildlife to cattle [16]. It is likely that local farming practices will impact on the bTB prevalence therefore it is important to identify local risk factors to *M. bovis* transmission [17].

Despite reports of isolation of *M. bovis* from livestock and wildlife, limited information is available in South Africa on the level of bTB knowledge and risk practices of livestock farming communities that influence the transmission of bTB to cattle and humans. The understanding of these practices will provide information for the development of informed grassroots programs that integrate the local and scientific knowledge towards bTB control in animal and zoonotic TB control in human populations living at the wildlife-livestock-human interface. The present study was therefore designed to understand the local knowledge on bTB among livestock farming communities and investigate the risk practices that were associated with bTB infection in cattle and in people co-existing with wildlife.

## Methodology

### Study area and population

A qualitative study based on focus group discussions (FGDs) was carried out in Big Five False Bay Municipality, uMkhanyakude District in the northern part of KwaZulu-Natal, South Africa. This community was part of a One Health investigation into the epidemiology of bTB at the wildlife-livestock-human interface. The Municipality is sparsely populated, and most of the population that occupy the north-eastern part are rural traditional communities with a cattle population that was estimated at 11 000 from a total of 456 owners (W. McCall personal Communication, 28 August 2017). The population is engaged primarily in crop-livestock farming and the main domestic animals are cattle and goats. The Municipality is surrounded by game and nature reserves such as St. Lucia (iSimangaliso), Hluhluwe/Imfolozi, Munyawana and Mkuze, which attract a high number of local and international tourists.

The study participants were purposively selected according to the inclusive criteria of being a household member of a farmer owning cattle that were tested for bTB in September 2016/ March 2017 at one of the four dip tanks in the area (Masakeni, Mpempe, Nibela and Nkomo) and owning either a bTB positive or negative herd [11]. A dip tank is a communal cattle handling facility where animals from several villages assemble weekly or once per fortnight for disease inspection and are also dipped in an insecticide plunge tank for external parasites control, primarily ticks.

The four-focus group categories that were selected from each dip tank included women that belong to households owning cattle, cattle keepers (male and female), heads of households and dip tank committee members. The group of adult female members of households was selected because in this socio-cultural context women are often solely responsible for the handling of food and food preparation for their families hence determining their consumption behavior. Cattle keepers were included because they are responsible for taking care of the animals (animal husbandry practices). These include young boys that are employees or members of the household (male/female). The heads of households are usually the decision-making group within the household that determine the movement, introduction and selection of animals for slaughter. Finally, the dip tank committee is a group of farmers selected by the community for each dip tank to assist the government animal health technicians in the management of dipping activities and reporting of animal diseases.

### Data collection

Fourteen focus group discussions (FGDs) were carried out in November and December 2017. In each FGD we examined the knowledge, awareness of bovine and zoonotic tuberculosis and the risk practices for bTB transmission to cattle, humans and wildlife among communal (rural) cattle farmers. A FGD guide was used and pretested in one FGD at Masakeni dip tank that consisted of dip tank committee members and the final topics examined include:

i. Local knowledge and awareness of bTB in terms of symptoms, transmission and control measures
ii. Food handling, preparation and consumption behaviors, cattle slaughtering procedures
iii. Livestock management practices in terms of cattle and wildlife interactions, cattle and cattle interactions and introduction of cattle into a herd.

A total of 150 people above 16 years were recruited by the local Animal Health technician into the study based on their various roles in the household and participation in the FGDs depended on their availability and or willingness. The participants for each group belonged to different households within the same dip tank, had similar socio-characteristics and were comfortable to discuss issues among themselves and the facilitator. FGDs were carried out at community halls that were centrally located and where village meetings or activities were commonly conducted.

The FGDs consisted of a facilitator, two observers and the preselected group of participants from the target community. The facilitator (TM), the first observer/note taker (PZ), and the principal investigator (PS) who was also a second observer in the discussion were trained in focus group interview principles and techniques by a qualified social science researcher (CV) from the Institute of Tropical Medicine, Belgium. The discussions lasted for about one hour, used the local language of isiZulu and were steered in a flexible and iterative manner. All the FGDs were led by the trained facilitator and the two observers, audio-recorded by digital voice recorder with the permission of the participants, transcribed and translated from local language into English. The facilitator and the observers were fluent in both English and isiZulu.

### Data management and analysis

All the discussions were transcribed into a Word document and these were cross checked and supervised by the principal investigator. The transcripts were read several times to get an overall understanding and to identify the main and salient themes until no new themes were found. The list of themes was used to code all the transcripts including the hand-written notes from the observers and the process was managed using NVivo Version 12 Pro^®^software. We examined the relationships between themes and sub-themes, patterns in the views expressed by the various groups or dip tanks in terms of differences or similarities. Each theme was described in detail and exemplary quotes were used to illustrate the meaning of the themes. The six main themes are represented in detail in the results section with quotations from the participants.

### Ethical consideration

The University of Pretoria-Faculty of Humanities-Research Ethics Committee approved the study (Reference:16394624/GWO170814HSA). At the commencement of each FGD the study was explained to the participants regarding the research purpose, FGD process, confidentiality and uses of the data. They could ask questions for clarity and thereafter they were given the consent form to read or it was read to them in the local language. A written consent was obtained from the participants to conduct the discussions and oral permission obtained to audio-record all the discussions.

## Results

The 14 FGDs involved the following groups from each dip tank; dip tank committee members, head of households, women from cattle owning households and cattle keepers from the 4 dip tanks; Masakeni, Nibela, Mpempe and Nkomo as shown in Table 1.

**Table 1:** Description of Focus Groups according to dip tank, number of participants and gender.

| Dip tank name | FGDs category | Number of participants | Gender |
| --- | --- | --- | --- |
| Masakeni | cattle keepers | 11 | male |
|  | women | 12 | female |
| Nkomo | head of households | 12 | male |
|  | cattle keepers | 11 | male & female |
|  | dip tank committee | 10 | male |
|  | women | 12 | female |
| Mpempe | head of households | 9 | male & female |
|  | cattle keepers | 12 | male |
|  | dip tank committee | 10 | male & female |
|  | women | 8 | female |
| Nibela | head of household | 12 | male & female |
|  | cattle keepers | 12 | male |
|  | dip tank committee | 10 | male |
|  | women | 9 | female |

The results are presented according to the themes that were identified from the analysis of the transcripts.

### Knowledge on bovine tuberculosis (bTB): symptoms, transmission and control measures

Most of the respondents indicated that they knew about bTB and considered it as one of the significant diseases of cattle found in their area. All the groups were able to discuss symptoms of the diseases. Knowledge on the transmission of bTB to cattle or humans was evident in all FGDs except the cattle keepers that were not sure of how this disease was transmitted to humans. Knowledge on bTB was shown by the ability of the participants to mention the correct clinical signs.

“*We have seen some of these from some of our animals. The animals cough a lot, I have heard the animals cough especially in the morning, the animal has difficulty in breathing making a loud sound, and it loses weight and has no appetite. That is all I know I do not know about the others”.* (Nibela dip tank, head of household, December 2017)

Most of the FGD participants were aware of possible transmission of zoonotic diseases from livestock to people through contact with infected animals or consumption of infected animal products, however they could not give specific examples of zoonotic diseases in livestock or wildlife.

“*[…] Let’s say there is a little child in the yard and the child approaches the cow as he or she (the child) is used to the cow and the cow is also used to this child, we might not see it at the time that the cow has a sickness. The child playing next to the cow can inhale whatever the cow coughs*”. (Masakeni dip tank, female member of household, December 2017)

The discussants from all FGDs groups, except the ones that included the women from cattle owning households, highlighted that transmission from infected cattle or wildlife to uninfected cattle was through direct contact or indirectly by grazing on contaminated pastures.

*“[…] The cows salivate uncontrollably and more than usual, so an infected cow can leave traces of saliva on the grass in which other cows can graze on and then become infected as well. TB can be transmitted because an infected cow is always coughing and lacks the means to cover itself when it does; the fact that it does not live in isolation means other cows and animals can easily be infected”.* (Mpempe dip tank, dip tank committee member, December 2017)

The participants were not conversant with bTB control methods instead mentioned that animal testing was needed since clinical signs could not be used for the diagnosis of the disease.

“*The problem with this disease that we are discussing is that it is only visible inside the animal and not outside. My biggest fear is that we may end up killing a lot of animals trying to figure out what is really bothering them. All I’m trying to say is that the knowledge that we have is not enough for us to make sound decisions”*. (Masakeni dip tank, cattle keeper, December 2017)

### Cattle slaughtering and meat inspection

The respondents from all the FGDs revealed that the selection of an animal for slaughter depended on the general health of the animal as observed by the owner, animal’s productivity (fertility), age and purpose i.e. for general consumption or for a traditional ceremony.

*“1. The reason for the selection of a cow is the size of the family and the number of guests expected. 2. Would be the overall health of the cow […]3. The cows are slaughtered or selected for slaughtering on a chronological basis where older cows stand a better chance of being selected than younger ones because of expected levels of yield from the young ones”.* (Mpempe dip tank head of household, December 2017)

The cattle keepers revealed that they did not wear protective clothing during slaughtering.

*“We handle the meat with these bare hands (shows hands), we don’t wear gloves for this. This is not a human corpse that has AIDS*”. (Mpempe dip tank cattle keeper, December 2017)

However, some respondents particularly from the dip tank committee members group had a different opinion after attending past educative meetings on bovine TB.

“*Back when we grew up that used to be the case, but now wearing sandals during slaughtering is no different than not wearing gloves. From all the meetings and information sessions we attend, the advice we get is ‘wear your protective clothing and gloves if you have’ and what is also different is that you have an animal inspected prior to it being slaughtered. In the olden days even, a sickly-looking animal was not spared […]”.* (Nkomo dip tank committee member, December 2017)

The respondents differed in their practices concerning the management of meat with spots. The cattle keepers revealed that in most cases the meat was not inspected and was consumed as quickly as possible once it was slaughtered:

“*Ours is different from the meat you get at the butchery where it is checked for abnormalities and illnesses. Once we have slaughtered the animal it is only correct for us to eat its meat”.* (Nkomo dip tank cattle keeper, December 2017)

The cattle keepers also highlighted that the consumption of uninspected meat was a common practice particularly during traditional ceremonies that could not be delayed or postponed.

*“The general procedure is that a cow must be checked before slaughtering. This isn’t followed because we want to eat meat during a ceremony. The way I see it, when it is said that we should check these cows before slaughter; what would happen if we find them to be sick and there is a huge possibility that many other are sick because they live together. Does this mean I must reschedule a settled date for an ancestral ceremony now that it was found that the cows are sick?”.* (Nibela dip tank cattle keepers, December 2017)

In contrast the FGDs that included cattle owners, dip tank committee members and the female household members indicated that they inspected the meat but that the occurrence of abnormalities was rare. In the case they were coming across unusual spots on organs such as the liver or lungs, they were taking the necessary precautions. These included discarding the affected organ by burying it in the ground, throwing it away or sending the organ to the animal health technician for testing but at the same time they proceeded to eat the rest of the meat that did not show visible spots.

“*We have a discussion amongst ourselves on what could possibly be wrong with the animal. After this we remove the affected area and continue with the healthy-looking parts”.* (Mpempe dip tank head of household, December 2017)

### Introduction of cattle into a herd by communal farmers

During the discussions it was clear that the circumstances that led to the introduction of cattle into the herds were common across all the dip tanks. These included the receiving of cattle as a ‘bride price’ as part of the marriage gifts from the groom’s family to the bride’s family, that is locally called “*lobola*”, when performing traditional marriage ceremonies and the commercial exchange of animals.

“*We do it for lobola or sometimes we sell amongst each other when we need money*”. (Nkomo dip, head of household, December 2017)

Other situations involved individuals that offered help to take care of their neighbor or relative’s cattle or when animals were exchanged to obtain a special bull for ‘breeding’ or a specific animal for a traditional ceremony as described below:

“*Maybe my neighbor needs a big cow for slaughter and to compensate for this exchange I may take two from his herd. You mentioned that there is a drought, another circumstance would be to move the cows to where conditions are better whether it is at a friend’s farm, neighbor or relative”.* (Masakeni dip tank cattle keeper, December 2017)

The animals that were used for compensation for a crime or paid as a fine were not mixed with the herd but were immediately slaughtered.

*“No, a cow for compensation does not stay long. It is slaughtered on the very same day and it doesn’t enter the yard because it is slaughtered at the gate on arrival. It does not stay long because it is an insult”.* (Mpempe dip tank, female member of household, December 2017)

### Criterion for accepting animals into the herd and the role of veterinary services

The criteria of acceptance of cattle into a herd was similar as explained in all the FGDs across the dip tanks and depended on the purpose of the introduction of the animals into the herd. For instance, cattle that were brought to the family to pay the bride price were not rejected due to social and cultural reasons.

“*Because everyone is happy during this time (Lobola and ultimately marriage), no one takes the time to thoroughly inspect the animals with a sober mind because of the happiness that the family may be feeling at the time. In a nutshell we do accept such cows knowingly or unknowingly*”. (Nibela dip tank cattle owner, December 2017)

*“[…] It is tradition, we live according to ancient rules and customs. When your prospective son-in-law comes to pay lobola and you refuse his cows you are guilty of a crime in this whole exchange. He (son-in-law) will never again be expected to pay Lobola because you refused to accept his cows initially. Customary law also agrees if he takes your daughter and goes on to live with her for free. He has the grounds and can state this at the tribal council that you refused to accept his cows when he was then willing. That is the primary reason why we accept cows. You are in part afraid of giving your son-in-law powers to walk away freely with your daughter while also demeaning your daughter’s character”.* (Mpempe dip tank cattle owner, December 2017)

The same principle of accepting the cattle without concern for their health status was also applied in circumstances whereby the transfer of animal was meant to help a neighbor or relative with animal care.

“*Another circumstance would be the exchange of cattle between owners when one has better grazing land and water conditions than the other. And during such an exchange it might be that my herd is infected and moving into uninfected herd and vice versa”.* (Nibela dip tank women, December 2017)

In response to the commercial exchange of animals all the groups concurred that in this scenario it was within their right to accept or reject an animal after inspecting its health status and if possible, involve their local state veterinary officers to grant the permission for movement of animals into the area.

“*What is better is the buying of cattle because I cannot buy something that does not satisfy me. But a cow that is to be given to me by someone else or one that needs help because of the drought, I would not turn back such person and his cow”*. (Masakeni dip tank head of household, November 2017)

### Cattle-to-cattle and cattle-to-wildlife interactions

Regardless of the group category, participants acknowledged that cattle-to-cattle contact from different herds was common in the area when their animals gather at the dip tank, communal watering points and nearby pastures.

“*Yes, our cattle interact with other cattle from herds different from ours. The interaction happens especially at grazing points and at the dip tank”.* (Nibela dip tank, head of household, December 2017)

Contact with wildlife was also noted when farmers were granted permission by the game park authorities to move their cattle into the game park for water and grazing. This information was provided by the respondents from FGDs that were mainly involved in the herding of animals i. e the cattle keepers, cattle owners and dip tank committee members. This situation was more generally the case of farmers from Nibela dip tank in our study.

“*There is a nearby wetland and a game park called iSimangaliso. Because of the difficulties posed by the drought, we have arranged with the park’s management to allow us to enter and let our cows graze and drink water within the park. And you would find our cows interacting with wildlife, especially buffalos, zebras and wildebeest drinking and grazing together”.* (Nibela dip tank committee members, December 2017)

Cattle to wildlife contact was also occurring through the broken-down fences surrounding some game parks in the area, allowing free movement of animals into and out of the game park.

*“The fence around the reserve is not properly erected to keep livestock out of the reserve and wild animals in. So, cattle love walking with zebras and with buffalos as well. This is even though buffalos are dangerous to the cow because they will kill it. Nevertheless, they do interact a lot”.* (Nkomo dip cattle keeper, December 2017)

However, one of the dip tanks (Masakeni) indicated that the fence that surrounded the game park in their area was not damaged and movement of animals into the game park was impossible.

“*In this area there is no wildlife-livestock interactions. No, they do not because wild animals from the game parks or lodges are well fenced in. It can be the cows and goats that interact with each other”.* (Masakeni dip tank cattle keepers, December 2017)

### Food preparation and consumption practices of communal farmers

Reference to the preparation and consumption of animal products for food, medicine and traditional practices was discussed with various groups.

Milk and meat were clearly stated as the main products that are obtained from the cattle.

*“It has to be meat, milk and amasi (soured milk)”*. (Mpempe dip tank women, December 2017)

Other animal products mentioned included cattle hide, horns, bile (for seasoning of meat and traditional rituals), gall bladder (for seasoning of meat and cultural wrist bands), urine (as medicine for a bad cough), cow dung (to neutralize poison) and blood which was mostly used following cultural practices.

“*Traditional healers and their trainees drink it (blood). Others drink the bile from the cow”*. (Nkomo dip tank women, December 2017)

The meat was prepared in various ways as illustrated by the responses given and whereby the meat was either boiled or grilled (“*braaiing”*).

“*My child, the meat is cut into different pieces, the portions that need grilling are cut out and portions that require to be boiled are also cut out. We leave it to boil in the pot till it is soft”.* (Masakeni dip tank women, December 2017)

The men, that mostly take part in the slaughtering of the cow, chose the meat portions for their *braaiing* or special stew and this determined the success of a traditional ceremony. Afterwards the rest of the meat is cooked by women for the whole family.

“*Methods of preparation vary with the kind of meat that is to be consumed. The norm is that there is meat called ‘stolen” meat. What I mean by this is that very small portions from the entire cow’s carcass are skillfully cut off to make a stew of all the portions cut off and the other half is flame-grilled (braai)”. (*Mpempe dip tank committee, December 2017)

When the participants were asked about their view on the consumption of undercooked meat the majority responded that it was the best and most preferred preparation, especially the liver, as expressed by a female participant:

*“Braai meat shouldn’t be overcooked my child, cooked so much that it becomes dry. The same applies to the liver. It must not dry up; it needs to have that ounce of blood on it because when it is well-done it becomes very hard”.* (Nkomo dip tank women, December 2017)

It was also clear that raw liver was consumed because they believed that it supplied nutrients.

*“Like those that eat a cow’s liver raw. Yes, raw. It is believed that there are more nutrients in it at this state than when it is cooked”.* (Mpempe dip tank committee member, December 2017)

Participants revealed that consumption of raw meat from the other organs of the cow was also done despite them knowing that this practice placed them at a risk for zoonotic diseases.

“*This would mean that the meat should not be eaten. The chances of us adhering to this condition are very minimal, because when we slaughter there is meat in which we consume sometimes raw or half-cooked. There are other parts in the cow’s intestine that are eaten raw”.* (Nibela dip tank, cattle keepers, December 2017)

A minority of the respondents disclosed that they were no longer eating undercooked meat although it was a difficult decision for them.

“*For health reasons, I no longer eat braai meat that is half cooked which oozes blood. It is difficult I will not lie, but I love my liver and as you would know it is not consumed well-done sometimes, it is eaten raw as this is believed that the liver in this state is very nutritious. Another meat consumed raw is the intestines, but a lot has changed since the discovery of diseases […]”*. (Nibela dip tank cattle owner, December 2017)

The discussants also acknowledged that in some traditional practices certain organs were meant for a specific group of people such as the chest for women, cow heels (trotters) and head of the cow for both young and elderly men and the liver which was given to the mother of a pregnant girl during marriage ceremonies or the elderly women in the family.

Milk is the other main product from the cattle that is consumed as either raw milk, boiled milk or as sour milk (“*aMasi”*).

“*We boil the milk but not always. When you are milking the cow you sometimes take straight shots into your mouth from the cow’s udder and it is nice and warm*”. (Nkomo dip tank cattle keeper, December 2017)

The preparation of “*aMasi”* was common to all groups and following traditional methods.

“*We first milk the cow and pour the milk into a traditional (calabash) or into 2-liter bottle of coke and it sits in a dry warm place maybe for three days and after that the “aMasi” should be ready*”. (Nkomo dip tank women, December 2017)

Milk was also used for traditional rituals for example during the cleansing ceremony of a widow, whereby the women used it to bath, and during funerals for the washing of hands.

*“Other families use it at funerals at the gate for the washing of hands when coming back from the gravesite. It is mixed with water”*. (Masakeni dip tank women, December 2017)

In some cases when a person had ingested poison or was constipated, milk was also used as a neutralizing substance or laxative respectively.

“*When you have ingested something poisonous or poison itself you mix the milk with some dung and then drink”*. (Masakeni dip tank, cattle keepers, December 2017)

## Discussion

A qualitative research approach was used to investigate the risk factors for bovine TB transmission to cattle to and humans using FGDs in the livestock-keeping community living at the wildlife-livestock-human interface. The awareness and knowledge of bovine TB in cattle and humans was also assessed using appropriate themes during the discussions. The purpose of the study was to document the community’s perception of bTB within their social and cultural context for the development of suitable interventions targeted towards the control of zoonotic TB. This is in line with the initiative by WHO/OIE/FAO/IUATLD (Road Map for Zoonotic Tuberculosis) to identify people at risk of zoonotic TB especially in sub-Saharan Africa in support of the WHO’s goal towards eradication of TB by 2050 [18].

Despite the participants being aware of zoonotic diseases they could not state specific examples of zoonotic diseases in livestock and wildlife. Contrastingly, a study in the Gauteng province of South Africa documented historical knowledge of zoonotic diseases amongst small-scale farmers and these recognized brucellosis as a zoonotic disease [19]. In other studies in sub-Saharan region cattle owners stated d rabies, anthrax, tuberculosis and brucellosis as major zoonotic diseases [20–22]. This lack of knowledge could be linked to the limited awareness campaigns on zoonoses, the absence of local information on zoonoses, inadequate communication between veterinary and human health professionals as described by Cripps [23]. Documentation from elsewhere in Africa indicates that the awareness, knowledge, attitude and perception of zoonoses determines the increase or decrease in zoonotic risk in livestock keeping communities and the general public [22,24].

Generally, the results showed a high awareness of bTB in cattle, the symptoms in cattle and mode of bTB transmission to cattle as well as to humans, although this was coupled with poor preventive practices. This is in contrast with other cattle keeping communities where the knowledge of bTB was generally found to be low as revealed by studies in Zambia, Tanzania and Ethiopia [25–27]. The high awareness in the community could possibly be attributed to the bTB activities associated with the research program on bTB conducted by our team which included the successful bTB information day and bTB testing of cattle at the dip tanks. Prior education on zoonotic diseases has been associated with a display of good knowledge of the disease by the cattle farming community as demonstrated in a study in Uganda [28].

Through the FGDs it was noted that most people are at least involved in one practice placing them at risk of bTB or other zoonotic diseases such as brucellosis. The risky practices that are characteristic of pastoral communities particularly in sub-Saharan Africa included consumption of raw (soured milk) or undercooked meat, poor handling practices and absence of trained veterinary personnel during slaughtering of animals [29– 31]. Most of the participants indicated that they consumed boiled milk, but this practice might have been overstated due to awareness by the participants of the accepted practice. Research has shown that people are inclined to display paradoxical behavior where one maybe aware but not apply precautionary measures [32]. Inadequate precautionary measures during production, processing, handling of animal products and slaughtering of cattle exposes individuals to bTB through direct contact with infected carcasses [8,20,33]. The poor practices might also be linked to the socio-economic status of the farmers whereby soured milk is readily available and cheap. Indeed, in contexts of fragile livelihoods, disease risk might not be people’s sole concern, or they might not have the resources to take up protection measures such as wearing protective clothing during slaughter [34]. The groups that were dominated by males, i.e. cattle owners, cattle keepers and dip tank committee members were at a greater risk of *M. bovis* infection than their female counterparts; as these were involved in unprotected practices during the slaughter of animals and consumption of undercooked meat during “*braaiing”*.

Consistent with other studies livestock keeping activities that were associated with bTB transmission to cattle involved uncontrolled movement of animals and introduction of animals into a herd without bTB pre-testing for social benefit, cultural or commercial purpose [7,15,35–37]. The free movement of cattle in communal farming within the village parameters is a common practice and movement to areas adjacent or into game reserves in search of pasture was attributed to the frequent drought conditions in the area. Livestock management practices that result in contact of uninfected herds with infected herds as well as livestock-wildlife contact have been observed in bTB endemic settings (South Africa and Zambia) and in the European context where occasional outbreaks have been reported [38–40].

The findings from this study were similar across all the dip tanks with minor differences concerning livestock-wildlife contact. Contact at the wildlife-livestock interface has been suggested to be caused by porous boundaries that promote sharing of resources that might result in the exchange of diseases between livestock and wildlife [41]. There were no extensive differences in animal husbandry, cultural practices, food consumption and handling behavior amongst the four different groups of participants since these groups belong to the same geographical location and tribe. However, there was a need to include all areas (dip tanks) with *M. bovis* infected herds for an in-depth analysis of risky practices.

Most of the opinions expressed by the participants in the four categories were similar with knowledge gaps or different views being expressed by the cattle keepers. Amongst the differences identified were the poor knowledge on bTB transmission to people, consumption of uninspected meat and the habit of drinking raw milk during milking, implying that the cattle keepers are more at risk of *M. bovis* infection. Several investigations have reported the potential risk of bTB transmission through drinking of raw milk and contact with infected cattle [42–44]. Poor practices regarding TB have been reported amongst high risk groups such as the cattle keepers and *M. bovis* transmission to people was confirmed in a study of livestock workers in Nigeria [29,45].

The study highlighted the influence of belief, habits and socio-cultural aspects on food processing (e.g. fermented milk), food consumption (undercooked or raw meat) and introduction of animals into a herd (lobola ‘bride price’). The consumption of raw or undercooked organs such as lymph nodes as influenced by socio-cultural practices has been previously reported in another study in South Africa [19]. Therefore, cultural issues related to drinking of raw blood, milk and meat (organs such as the liver and intestines); selection of animals for slaughter during traditional ceremonies and acceptance of animals into a herd without bTB pre-testing might be difficult to alter. Instead farmers should be informed about potential consequences of certain practices in the spread of bTB in cattle and people. Cultural practices are recognized as impediments in *M. bovis* control strategies in developing countries, consequently 10-15% of human TB is potentially caused by this pathogen [46]. Thus, policy makers need to be familiar with the cultural practices associated with the cattle owners’ actions that influence bTB control to be able to design effective preventive solutions.

Most of the data on bTB risk practices in previous studies was obtained using quantitative methods only or mixed with qualitative methods, whilst in this study a qualitative method of FGDs was applied. FGDs provide a platform for participants to easily discuss their beliefs within the group atmosphere than individual interviews [47]. Using the different categories according to the people’s role in the community/households allowed the participants to be comfortable and freely express their opinions. The advantage of using FGDs is that the researcher can reach many people on one goal, explore people’s knowledge, experiences and collect a large amount of data within a short period of time [48]. The limitation of this method is that the researcher has no control over the information that is generated during the discussion and it’s not the best way to obtain responses on sensitive issues [49]. More participant observations would have triangulated some data regarding practices (e.g. milk consumption), hence avoiding bias raised during the discussions.

Using disease risk as a departure point in One Health studies of zoonoses is very pertinent as reducing risk has a real public health, economic and social rationales. However, it is very important to avoid that people’s cultural logics or social practices are cast in negative terms, as ignorance or superstition, or behaviors that exacerbate risk and require changing [34]. Furthermore, how risk is perceived by livestock owners or by animal and human health workers are not necessarily the same [50]. The endeavor of addressing risky practices is complex and cannot involves a simple linear process of social engineering because knowledge alone does not drive behavior change. Derived from the social and psychological sciences, Kelly and Barker (2016) proposed to start with the behavior, identify who is behaving and where, and working backwards using regressive inference (understand the preceding conditions of the specifics) [51]. It is a much more profitable avenue for developing interventions instead of predictive single causal models such as the model of Theory of Planned Behavior which is elaborated on rational assessment alone based on economic utility theory [51,52].

## Conclusion

Our qualitative study allowed us to inform about a nuanced understanding of how people experience and indeed conceptualize risk within specific socio-cultural practices and a wider web of structural factors such as the intersection of poverty and gendered divisions of labor affecting risk of bTB in an endemic area at the wildlife-livestock-human interface in northern Kwa-Zulu Natal, South Africa. These findings are keys for the elaboration of appropriate public health policy and related education campaigns. The strengths of using a qualitative approach is its attention to complexity, questioning the familiar, helping with language and translation, reconfiguring boundaries (e.g. human-animal relations) to create novel frameworks and being reflective [53]. The participants displayed good knowledge of bTB in cattle and its transmission to humans and cattle. However, the perceived risks to humans and cattle were not translated into protective practices as these were influenced by socio-cultural aspects and economic value of livestock keeping in the community. Finally, concerted effort is required from all stakeholders using the One Health strategy for the implementation of an all-encompassing disease control program as control of zoonotic TB transmission to humans is ultimately linked to the control of bTB in cattle.

## Recommendations

A community-based animal health delivery system that provide basic instructions on practices that reduce risk of disease transmission to cattle and people through increased awareness campaigns is suggested. We suggest community involvement in the planning of appropriate educational programs that consider human-animal relations, social, cultural and economic reality of the community. These programs should involve the community leaders (tribal leaders) that are respected by the community, health caregivers and local veterinary personnel. In addition, targeted educational programs would benefit risk groups, for example cattle keepers to improve their perception towards zoonotic TB since the group is actively involved in livestock keeping activities.

## Acknowledgements

The authors are indebted to the team of investigators involved in the study Tebogo Mogoru (TM) who was the facilitator and Philani Zulu (PZ) first observer during the FGDs. We acknowledge the veterinary staff at Big 5 False Bay Municipality offices especially Mr. Simon Mkwamubi (Animal Health Technician) for their contribution to the success of the study. We also appreciate the local tribal authorities for granting us the permission and the farmers who agreed to participate in the study.

